# NRF2 drug repurposing using a question-answer artificial intelligence system

**DOI:** 10.1101/594622

**Authors:** Michel-Edwar Mickael, Marta Pajares, Ioana Enache, Gina Manda, Antonio Cuadrado

**Affiliations:** Victor Babes institute, Bucharest, Romania; PM forskning center, Stockholm, Sweden; Autonomous University of Madrid

**Keywords:** Computational immunology, NRF2, text mining, QA, ranking

## Abstract

Drug repurposing represents an innovative approach to reduce the drug development timeline. Text mining using artificial intelligence methods offers great potential in the context of drug repurposing. Here, we present a question-answer artificial intelligence (QAAI) system that is capable of repurposing drug compounds. Our system employs a Google semantic AI universal encoder to compute the sentence embedding of an imposed text question in relation to publications stored in our RedBrain JSON database. Sentences similarity is calculated using a sorting function to identify drug compounds. We demonstrate our system’s ability to predict new indications for already existing drugs. Activation of the NRF2 pathway seems critical for enhancing several diseases prognosis. We experimentally validated the prediction for the lipoxygenase inhibitor drug zileuton as a modulator of the NRF2 pathway in vitro, with potential applications to reduce macrophage M1 phenotype and ROS production. This novel computational method provides a new approach to reposition of known drugs in order to treat neurodegenerative diseases. Github for the database and the code can be downloaded from https://gist.github.com/micheledw/5a165b44345d45105d715340b88c756b

## Introduction

### Problems with current drug repositioning techniques

Compound identification is one of the main bottlenecks of drug repurposing (repositioning). Drug repurposing constitutes a paradigm shift from traditional drug development approaches [1], since traditional drug discovery workflows suffer from high cost and long delays. Developing a single new drug can cost over one billion dollars [2]. Reports estimate the duration of a traditional drug discovery cycle to be fifteen years [3]. The largest segment of time and expenses of this cycle is allocated to early development, with more than 90% of drugs failing to move beyond the first stage [4]. Repurposing of drugs that have been approved for human treatment diminishes the expenses linked with the early stages of drug development. Furthermore, this approach can reduce the delay faced by therapeutic indications. Successful examples of drug repositioning include sildenafil for pulmonary hypertension, thalidomide for erythema nodosum leprosum, and retinoic acid for promyelocytic leukaemia [4]. Compound identification together with compound acquisition, development, and post-market safety monitoring constitute the main phases of the drug repurposing workflow [5]. In addition to its long duration, which lasts around two years, compound identification phase is also taunted by low success rates. Thus, enhancing the efficiency of compound identification is critical for improving drug discovery research.

### AI reduces the resources needed for drug repurposing

Biological question-answer systems employing AI could reduce the resources needed for the compound identification phase. AI has revolutionized text mining in the context of question-answer systems [6]. Manual extraction of data is biased and time-consuming [7]. Automatic text mining can accelerate the processes of knowledge formation and hypothesis building [8]. However, drug terms and vocabulary have made building tools for this specific research area challenging. To solve this problem, machine-learning methods could be used to identify these terms in a context-dependent manner [8]. This approach was applied to infer protein-protein networks and to identify genes associated with diseases such as breast cancer [9]. Unfortunately, this method requires text standardization processes (e.g. stemming, lemmatization) [10]. Employing semantic similarity techniques have the potential to solve this problem. Nevertheless, current models suffer from low accuracy To tackle this problem, an AI approach was recently proposed [6]. Using a universal sentence encoder, researchers were capable of encoding text into high dimensional vectors (HDV) using a syntactic composition function [11]. HDV can be used for text classification, semantic similarity, clustering and other natural language tasks with considerably higher accuracy [6]. However, the full potential of applying AI in QA systems for drug discovery has not been achieved yet.

### NRF2 influence macrophages in neurodegenerative diseases

NRF2 pharmacological activation could play a vital role in regulating ROS in macrophages during neurodegenerative diseases. In neurodegenerative diseases, ROS are capable of producing membrane damage, changes in the inner proteins affecting their structure and function, lipids denaturation, and structural damage to DNA in the brain [12] [13]. ROS also play a major role in gradual deterioration of macrophages functional characteristics in neurodegenerative diseases [14] [15]. Oxidative imbalance produce reactive carbonyls that alter the ECM environment of macrophages decreasing their phagocytic activity towards apoptotic cells [16]. Furthermore, oxidative and carbonyl stress inhibit the activity of the transcriptional co-repressor HDAC-2 which under normoxic conditions helps to suppress pro-inflammatory gene expression [16]. The CNS is equipped with a repertoire of endogenous antioxidant enzymes, which are regulated by the transcription factor NRF2 [17]. Under normal unstressed conditions, NRF2 is bound to KEAP1[18]. Under conditions of oxidative stress by either reactive electrophiles, toxins or ARE inducers, the interaction between NRF2 and KEAP1 is disrupted. NRF2 translocates to the nucleus, where it binds to Smaf proteins [13]. This process increases the transcription rate of antioxidant response element [13]. Interestingly, NRF2 was shown to be up-regulated in multiple sclerosis plaques and primarily expressed in macrophages [17]. Moreover, NRF2 suppresses lipopolysaccharide-mediated macrophage inflammatory response by blocking IL-6 and IL-1β transcription, in EAE mouse models [19]. It was also demonstrated that, NRF2-mediated antioxidant gene expression can reduce the macrophage M1 phenotype and ROS production [20]. Using NRF2 activators has become a potential therapeutic strategy for numerous neurodegenerative diseases [20] [21]. However, the number of NRF2 activators applied in clinics is still small. Tecfidera (dimethyl fumarate), a potent NRF2 activator, has been approved for the treatment of multiple sclerosis but long-term use of this drug can cause resistance and other side effects [21] [22]. Nardochinoid C was reported to inhibit inflammation and oxidative stress in lipopolysaccharide-stimulated macrophages. However, its effect on neurodegenerative diseases is not yet known. Therefore, evidence indicates that the discovery of new and safer NRF2 activator for clinical use in neurodegenerative disease has become an important task in drug discovery [20].

### Paper objective

The objective of this study is to identify putative chemical compounds able to activate NRF2 in macrophages using our QAAI system. Using this approach, we searched 100 previously published PubMed publications for compounds that could affect oxidative stress in chronic diseases. Our approach highlighted twenty-five compounds. From these 25 compounds, we evaluated the ability of the lipoxygenase inhibitor Zileuton to cross the blood brain barrier. Afterwards, we investigated its ability to activate NRF2 activation in the macrophage cell line RAW26.7. Here we present evidence that our strategy can uncover new uses for FDA-approved drugs.

### Methods

The question-answer pipeline could be divided into five stages (Figure 1). The first stage aims to build a JSON database. Secondly, the sentence embedding was calculated for each publication together with the imposed question. Thirdly, all putative answers were ranked using an optimized sorter function and the highest answer presented. After that, we performed an *in silico* simulation to examine the ability of the identified drugs to cross the blood-brain barrier. Finally, we experimentally validated one of the drug targets identified by our QAAI system.

#### (i) Building the data base

Our database was built in JSON format using publicly available literature. We downloaded 100 files in PDF format from PubMed with the query term “chronic diseases” “drugs” “oxidative stress”. Text was then extracted from each of these files utilizing Tika python library (v 1.19) using “parser.from_file” function to parse each file [23]. After that, the Natural Language Processing (NLP) python module was employed to perform sentence tokenization into unique sentences with the tokenizer “nltk.data.load english.pickle” [24] [25]. All tokenized sentences were then written into the database using json.dumps function in JSON module [26][27].

#### (ii) Sentence similarity encoder

We employed Google universal encoder to calculate sentence embedding in Tensorflow. Google universal encoder uses a deep averaging network (DAN)[28]as its composition function. The primary advantage of the DAN encoder is that compute time is linear to the length of the input sequence [6], [19]. By this approach, input embedding for words and bi-grams are first averaged together and then passed through a feedforward deep neural network to calculate sentence embedding [18]. Overall, the encoder takes as input a lowercased Penn Treebank (PTB) tokenized string and outputs a 512-dimensional vector as the sentence embedding. The code was run in the Jupyter notebook on google Colab [11][29].

#### (iii)Evaluation of the best sorting algorithm

We benchmarked several functions to identify the most accurate correlation technique. We analyzed the answers generated for ten manually benchmarked questions using the built database. The correlation between the questions embedding and each sentence embedding was calculated using two different approaches (i) Inner product, (ii) Gromov–Hausdorff (supplementary file 1). The results were then sorted and the highest answer was assumed to be the most relevant. Then, for each of the ten questions, the values of true positive, true negative, false positive and false negative were calculated. The true positive is defined as the ability of the workflow to locate a sentence in the document that contains the name of a compound. Finally, we computed the f-score to evaluate our different correlation approaches (supplementary file 1).

### *In silico* prediction of blood brain barrier diffusion

The ability to cross the blood-brain barrier (BBB) is the main obstacle facing neurodegenerative treatment drug development. To tackle this problem, we utilized BBB predictor (http://www.cbligand.org/BBB) to investigate the ability of zileuton to cross the blood-brain barrier. First, zileuton chemical structure was downloaded from DrugBank (https://www.drugbank.ca/drugs/DB00744) in PDB format. Then, BBB predictor was employed using two different algorithms (i) support vector machine (SVM) and (ii) LiCABEDS. To ensure consistency, we employed four types of fingerprints (e.g. MACCS, Openbabel, Molprint 2D and PubChem). Finally, the score of the BBB crossing probability was calculated using default settings.

### Experimental validation

RAW264.7 cells were grown in Roswell Park Memorial Institute (RPMI) supplemented with 10% fetal bovine serum (HyClone, CH30160.03) and 80 μg/ml gentamicin (Laboratorios Normon, 763011.1). Cells treated with Zileuton (Sigma-Aldrich; Z4277) in the absence of serum. After the indicated times, cells were lysed in lysis buffer (50 mM Tris-HCl pH 7,5, 400 mM NaCl, 1mM EDTA, 1 mM EGTA, 1% SDS, 1 mM PMSF and 1 µg/mL leupeptin), and samples were heated at 95°C for 15 min, sonicated and precleared by centrifugation. Protein was quantified with DC(tm) Protein Assay (Bio-Rad). Primary antibodies were the following: anti-NRF2 (homemade; 1:5000), anti-HMOX1 (Enzo life sciences OSA110; 1:2000), anti-ACTB (Santa Cruz sc-1616; 1:5000) and anti-LAMINB (Santa Cruz sc-6217; 1:5000). Membranes were analyzed using the appropriate peroxidase-conjugated secondary antibodies (anti-mouse and anti-rabbit from GE Healthcare UK Limited, NA931V and NA934V, and anti-goat from Santa Cruz, sc-2020). Proteins were detected by enhanced chemiluminescence (GE Healthcare, RPN2232).

## Results

Our results point to zileuton as a putative NRF2 modulator in neurodegenerative autoimmune diseases. We built a JSON database of oxidative stress-related publications that carried the words “oxidative stress”, ‘chronic diseases” and “drug”. Our workflow computed the embedding for each sentence stored in the database using a Google encoder deeply averaged network. Following that, we calculated sentence similarity between the imposed question and every stored sentence in the database. Then, we used two different functions to calculate the similarity scores. Out of 25 compounds identified by our algorithm, we focused on zileuton, as the most relevant candidate. Moreover, we investigated its ability to cross the blood-brain barrier *in silico* and validated its ability to activate NRF2 *in vitro*. Our strategy could reduce the time and the costs for compound identification in the field of drug repurposing to tackle neurodegenerative diseases.

### Model performance

It seems that the best algorithm for calculating the sentence similarity is using the inner product algorithm. The embedding of each sentence was calculated (Figure 2). After that, the sentence similarity score was computed (Figure 3). For each method, we calculated the accuracy scores (Figure 4). The inner product method f-score (0.67) was higher than that of the Gromov–Hausdorff (0.18).

### Results highlighted by the AI program

The artificial intelligence (AI) workflow input was the question “what are the drugs that affect oxidative stress in chronic diseases?”. The workflow searched 100 documents for relevant answers. Each sentence of every document as well as the imposed question was converted to a unique numerical vector using an AI encoder. Then, the correlation between each vector and the question vector was computed using the inner product function. For every document, the answer was assigned to be the sentence with the highest score (List 1).

### Blood brain barrier passive diffusion ability

The results suggest that the BBB is preamble to zileuton. Using the AdaBoost algorithm, the score of BBB crossing was 10.985± 6.195189. Utilizing the SVM (which employs a different scale) generated a score of 0.3025±0.2396838. This indicates that zileuton (Figure 5) has the ability to cross the BBB (Table 1, Figure 6).

**Table 1.** Summary of probabilities of zileuton crossing the BBB.

| Test | Threshold | Score |
| --- | --- | --- |
| AdaBoost_MACCSFP | 0 | 4.10 |
| AdaBoost_OpenbabelFP2 | 0 | 17.6 |
| AdaBoost_Molprint2DFP | 0 | 7.66 |
| AdaBoost_PubChemFP | 0 | 14.58 |
| SVM_MACCSFP | 0.02 | 0.027 |
| SVM_OpenbabelFP2 | 0 | 0.344 |
| SVM_Molprint2DFP | 0 | 0.603 |
| SVM_PubChemFP | 0 | 0.236 |

### Experimental validation results

Zileuton increases NRF2 levels (Figure 7). RAW264.7 cells treated with zileuton (10 µM, 16 h) in the absence of serum showed increased NRF2 levels compared to vehicle-treated control cells. It is worth noting that we also observed an increase in HMOX1 protein levels. As expected, the increase in HOMOX1 was larger than NRF2. Heme oxygenase-1 (HMOX1) is regulated by NRF2. Presumably, NRF2 levels increase first, and it has to translocate into the nucleus, induce HMOX1 transcription which has to be translated.

**List 1.**
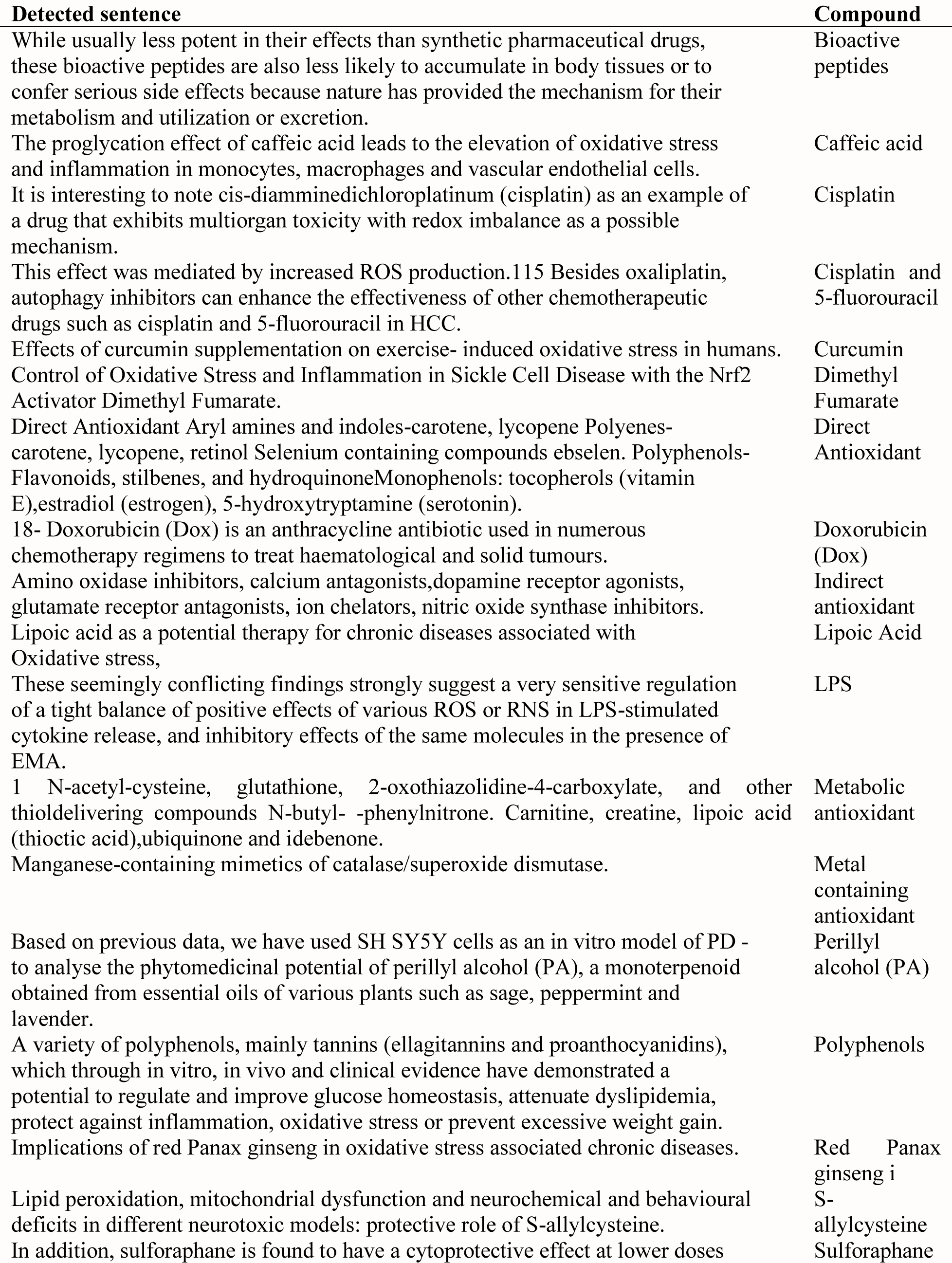
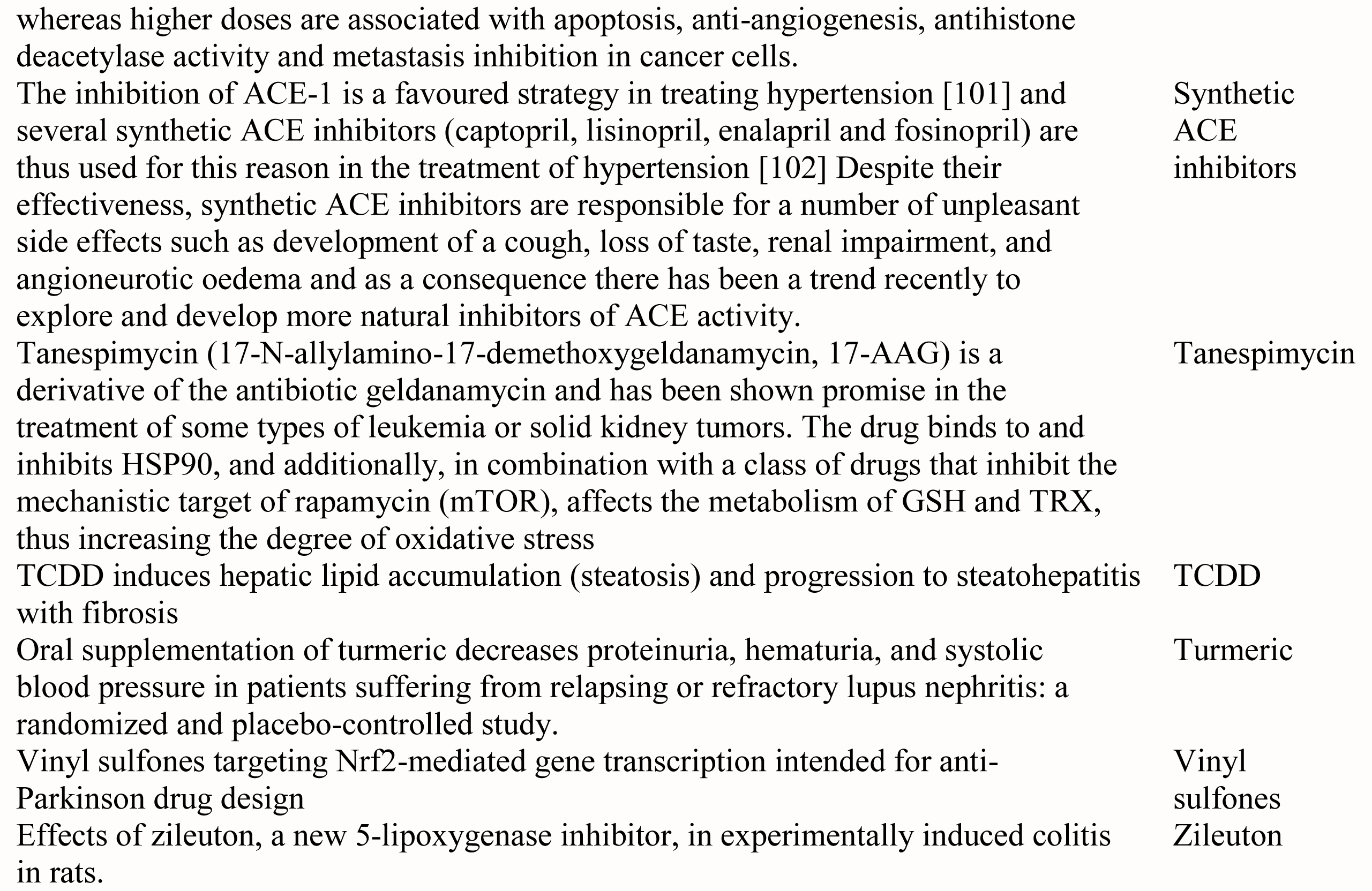
Results of the question “what are the drugs that effect oxidative stress in chronic diseases?”.

## Discussion

We focused on NRF2 activation drug repurposing using an AI approach in Google Colab environment. In biomedical applications, semantic similarity has become a valuable tool for analyzing the results in gene clustering, gene expression and disease gene prioritization [30]. In addition, the use of sentence similarity include estimating relatedness between search engine queries [2] and generating keywords for search engine advertising are active areas of research [3]. Our approach further extends these areas to make use of hundreds of drugs already approved for human usage. Our pipeline first calculates sentence embedding using a deep averaging network encoder. Then, we calculate sentence similarity between the posed question and the available dataset. Our system identified zileuton as a putative compound to tackle neuroinflammation in neurodegenerative diseases. Interestingly, we predicted its ability to cross the blood-brain barrier by an *in silico* method. Moreover, we validated its ability to induce NRF2 and its target HMOX1 levels in a macrophage cell line. Our approach seems capable of opening more opportunities for drugs repurposing.

### AI in text mining approach

Our system provides a context-aware approach for drug repurposing. The vast pharmacological knowledge available in the literature has made it increasingly feasible to employ text mining drug indications approaches. Swanson’s ABC links two concepts using a common relationship [31]. The clinically verified drug repurposing of fish oil to treat Raynaud’s syndrome was achieved using this approach[32]. However, approaches based on concept co-occurrence within abstracts generate a high percentage of false positives hypotheses[33]. Another alternative is network-based approaches. DrugMap Central[34] is a network-based approach that uses information on chemical structures, drug targets, and signaling pathways to de novo alternative drug indications. However, this approach is time-consuming and also generates many hypotheses. Tari et al [33] employed a parse tree which is an ordered, rooted tree that represents the syntactic structure of a string according to a context-free grammar. In addition to being context-free, their method also is ontology-based. Our method, however, eliminates the need for ontology through a context-aware AI system.

### Sentence similarity methods

Using the inner products of the embedding vectors proved to be the most accurate approach (Figure 4). There is a wide range of methods for calculating the similarity in meaning between two sentences including (i) Baseline, (ii) Word Mover’s Distance (iii) Smooth Inverse Frequency and (iv) AI encoders. Baseline method calculates the average of the word embedding of all words in the two sentences and then calculates the cosine between the resulting embedding[35]. This method lacks consistency as well as accuracy[35]. It also gives weight to irrelevant words[35]. Word Mover’s Distance measures the minimum distance that the words in one text need to “travel” to reach the words in the other text[36]. However, this method is slow. Smooth Inverse Frequency method tackles the problem of irrelevant words by using a weighted average of the word embedding [37]. It also removes common components by calculating the PCA for every sentence[37]. However, PCA is computationally complex and subject to random fluctuation. Here, we used a pre-trained Google universal Encoder which has proven to be more accurate, less time consuming but more memory intensive[6]. In order to optimize its function, we added a correlation unit that takes the sentence embedding as its input and then calculates sentence similarity using different correlation algorithms. The correlation method that possess the highest level accuracy seems to be the inner product approach. The reason behind could be that the Gromov–Hausdorff distance is essentially intractable as it involves the solution of an NP-hard optimization problem [38]. Thus, we employed the directed Hausdorff distance as an approximation to the Gromov–Hausdorff. However, the directed Hausdorff distance accuracy could be affected by the nonlinear nature of the embedded matrices.

### Zileuton modulation of NRF2

Our research repurposes zileuton as pro-inflammatory enhancer by activating NRF2 in macrophages. Oxidative stress is known to increase the levels of free arachidonic acid [39]. Free AA can be converted to bioactive eicosanoids through the cyclooxygenase (COX), lipoxygenase (LOX) or P-450 epoxygenase pathways[40]. LOX enzymes (5-LO, 12-LO, 15-LO) catalyse the formation of LTs, 12(S)hydroperoxyeicosatetraenoic acids and lipoxins (LXs), respectively [41]. COX isozymes (constitutive COX-1 and inducible COX-2) catalyze the formation of PGH2 [42]. PGH2 is converted by cell-specific PG synthases to active prostanoids (including PGE2, PGF2a, PGI2 and TXA2) [41]. 5-lipoxygenase is found throughout the central nervous system, in both neuron and glia [43]. 5-lipoxygenase is also active mainly in myeloid cells, such as macrophages [44]. LTs are made predominately by inflammatory cells such as activated macrophages [45]. Zileuton is an active inhibitor of 5-lipoxygenase, and thus inhibits leukotrienes formation. Treatment with, zileuton, at an early stage of the development of the AD-like phenotype delays cognition impairments, reduces amyloid beta (Aβ) levels, and tau phosphorylation in mouse models of AD[46]. Zileuton treatment of AD-like phenotype of the 3xTg mouse model of AD starting at 12 months of age restored mice cognitive abilities and reduced Aβ deposition as well as tau phosphorylation [46]. It was also shown that blocking of 5-lipoxygenase with zileuton delayed the onset and reduced the cumulative severity of EAE mice[47]. Zileuton also possesses the ability to suppress prostaglandin biosynthesis by inhibition of arachidonic acid release in macrophages [48]. In confirmation with these studies [46], we observed the ability of zileuton to activate NRF2, probably through the prostaglandin pathway (Figure 8). We also demonstrated an increase in the downstream target of NRF2, HMOX1. HMOX1 possess anti-inflammatory properties through up-regulation of interleukin 10 (IL-10) and interleukin 1 receptor antagonist (IL-1RA) expression[49]. This could further skew macrophages polarization towards an M2-like phenotype [50]. Our investigation predicted that the BBB is preamble to zileuton. This suggests that increasing the activity of NRF2in macrophages using zileuton could have therapeutic applications in neurodegenerative diseases.

### Limitations and future development

Although our approach is innovative, it still suffers from several limitations. On the computational side, our method employed google recurrent network encoder using the transformer architecture. However, we have not compared its performance with other available encoders such as inferSent which is a pre-trained encoder that was developed by Facebook Research[51]. We did not apply any mathematical optimization techniques to identify the lowest possible Hausdorff distance. In future versions of QAAI, we will also add a server and a platform to expand the usability of the system. From an experimental point of view, we have not investigated the effect of zileuton on microglia which plays a critical role in neurodegenerative diseases. It will also be interesting to validate the ability of zileuton to cross the BBB either by passive diffusion or active transport.

## Conclusion

We developed a systematic computational method to predict novel therapeutic indications through text mining of publically available publications. We demonstrated the ability of our QAAI system to predict novel therapeutic relationships, one of which is experimentally validated *in vitro*.

## Supporting information

Main suppl-table-mkr

## Acknowledgement

We would like to acknowledge Dr Macrious Abraham for his ideas and advice regarding the text mining part.

---

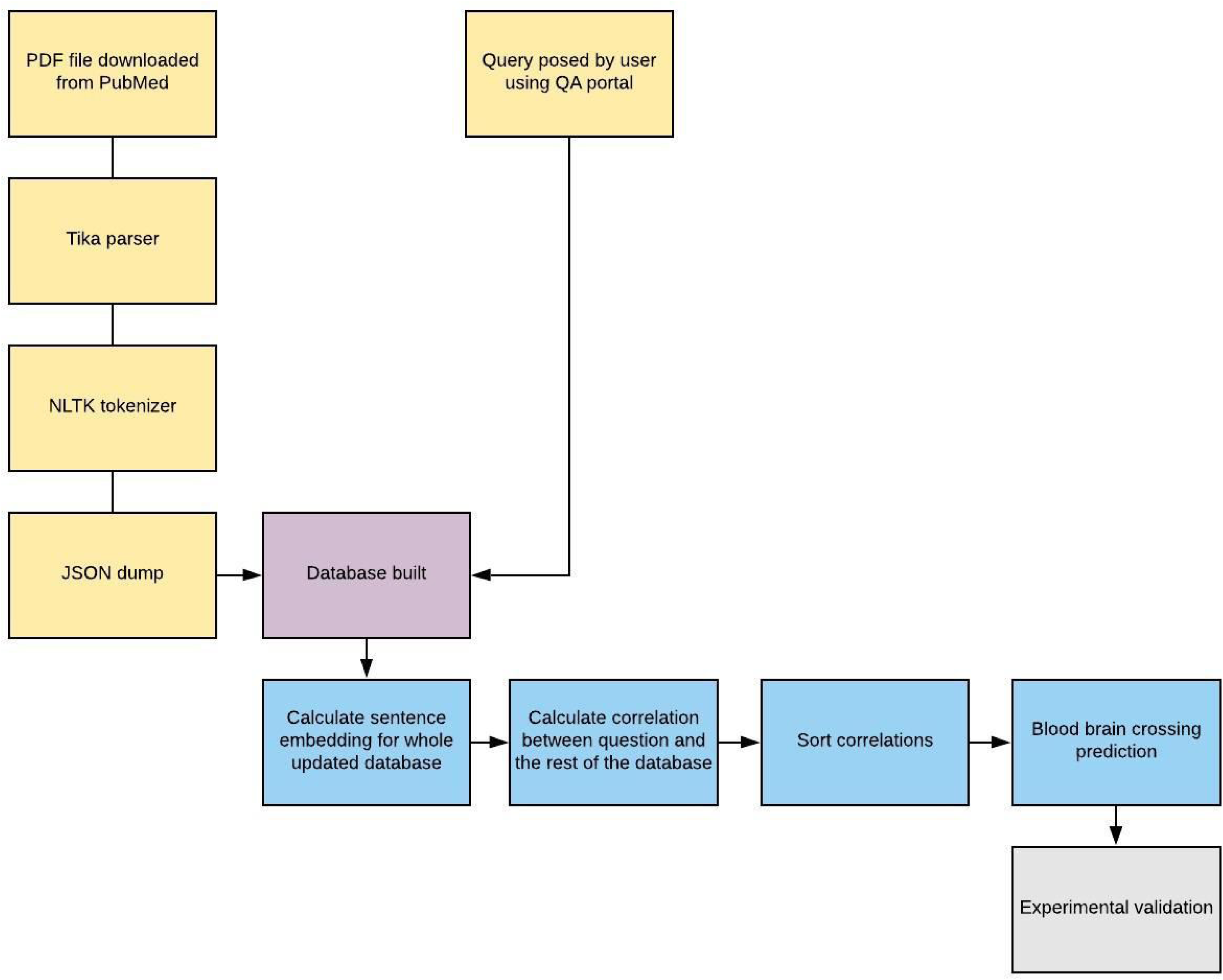

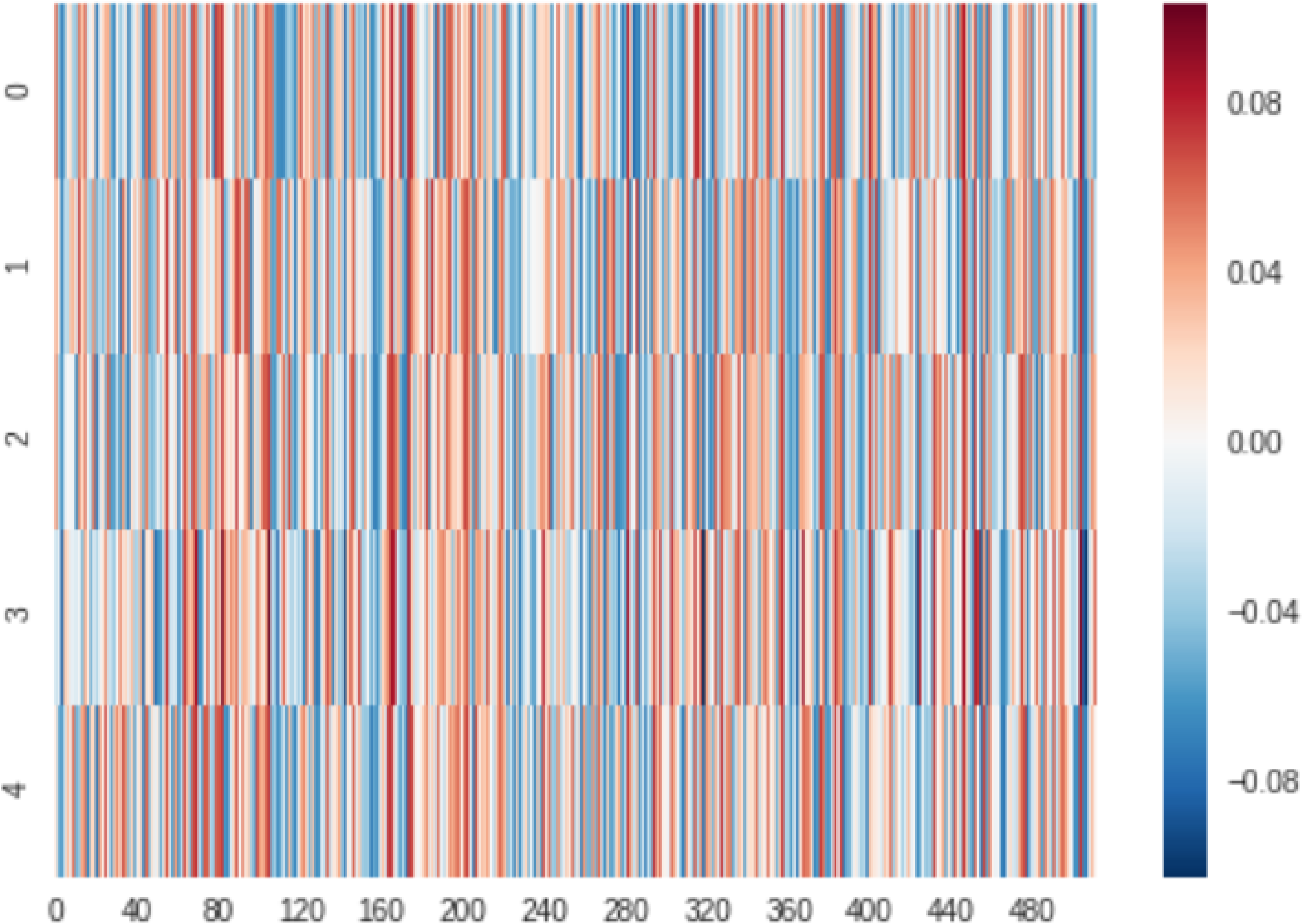

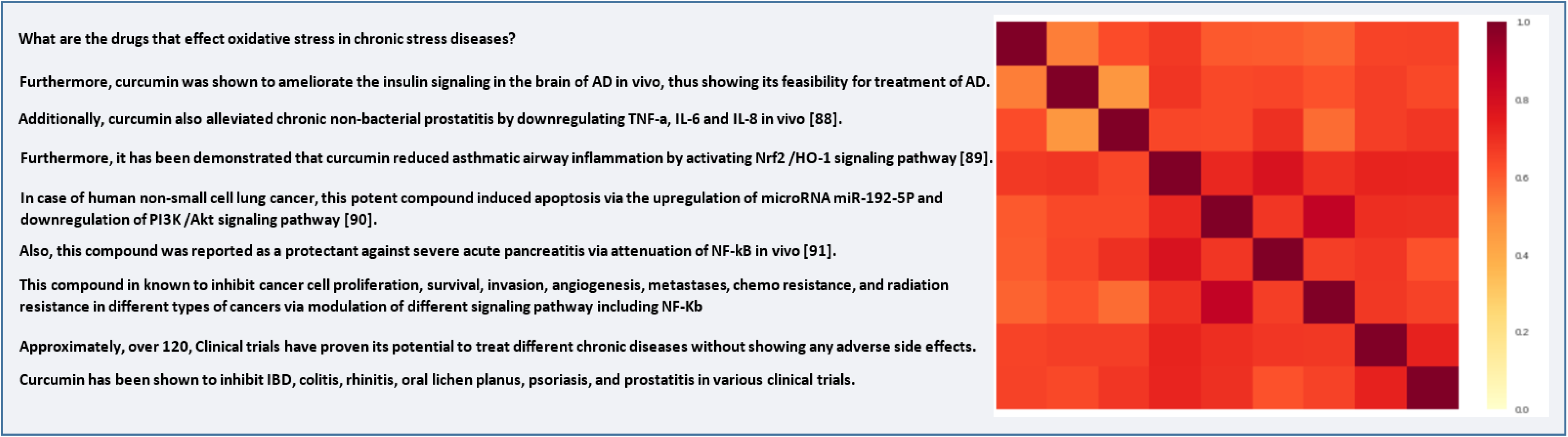

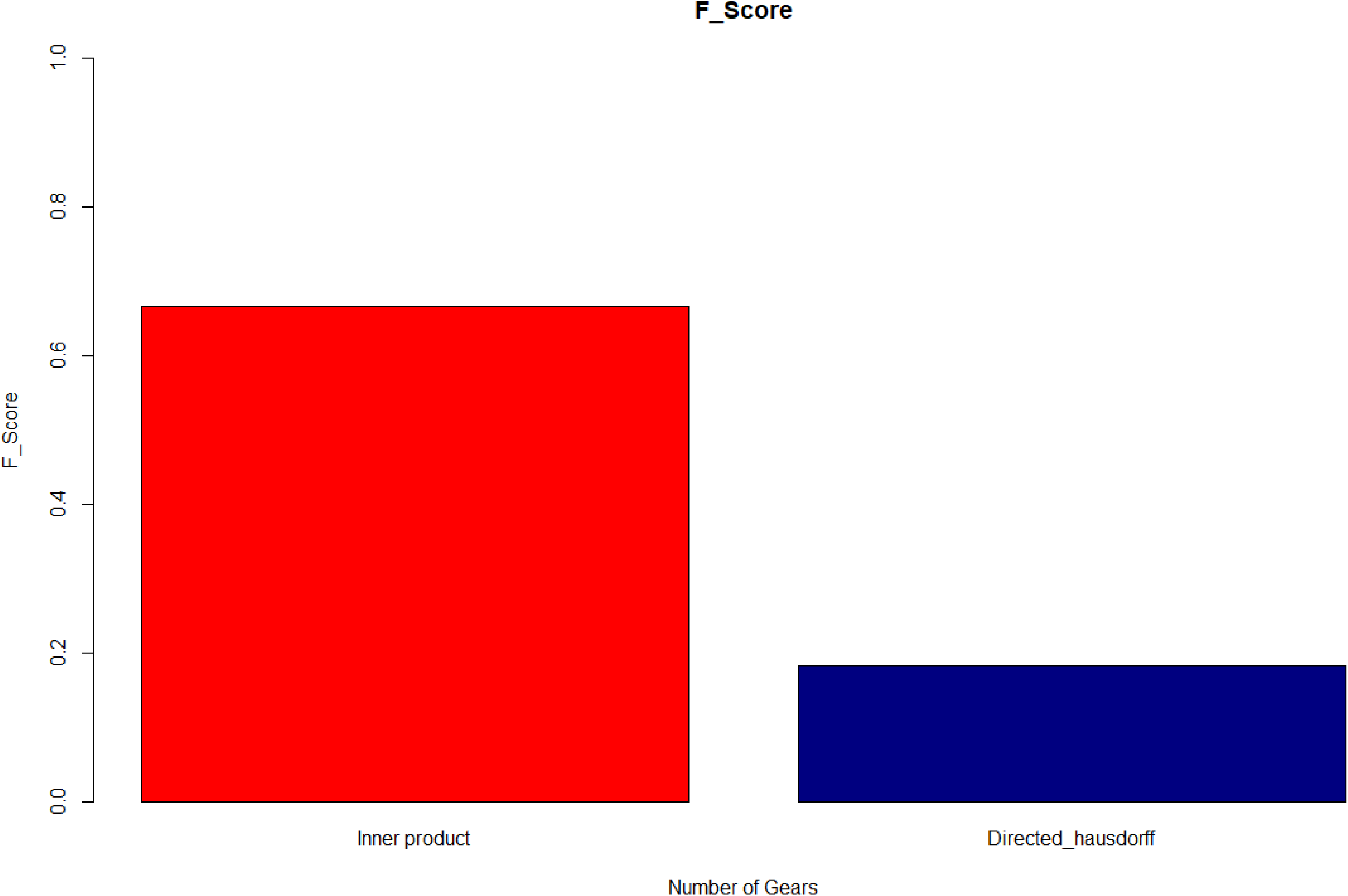

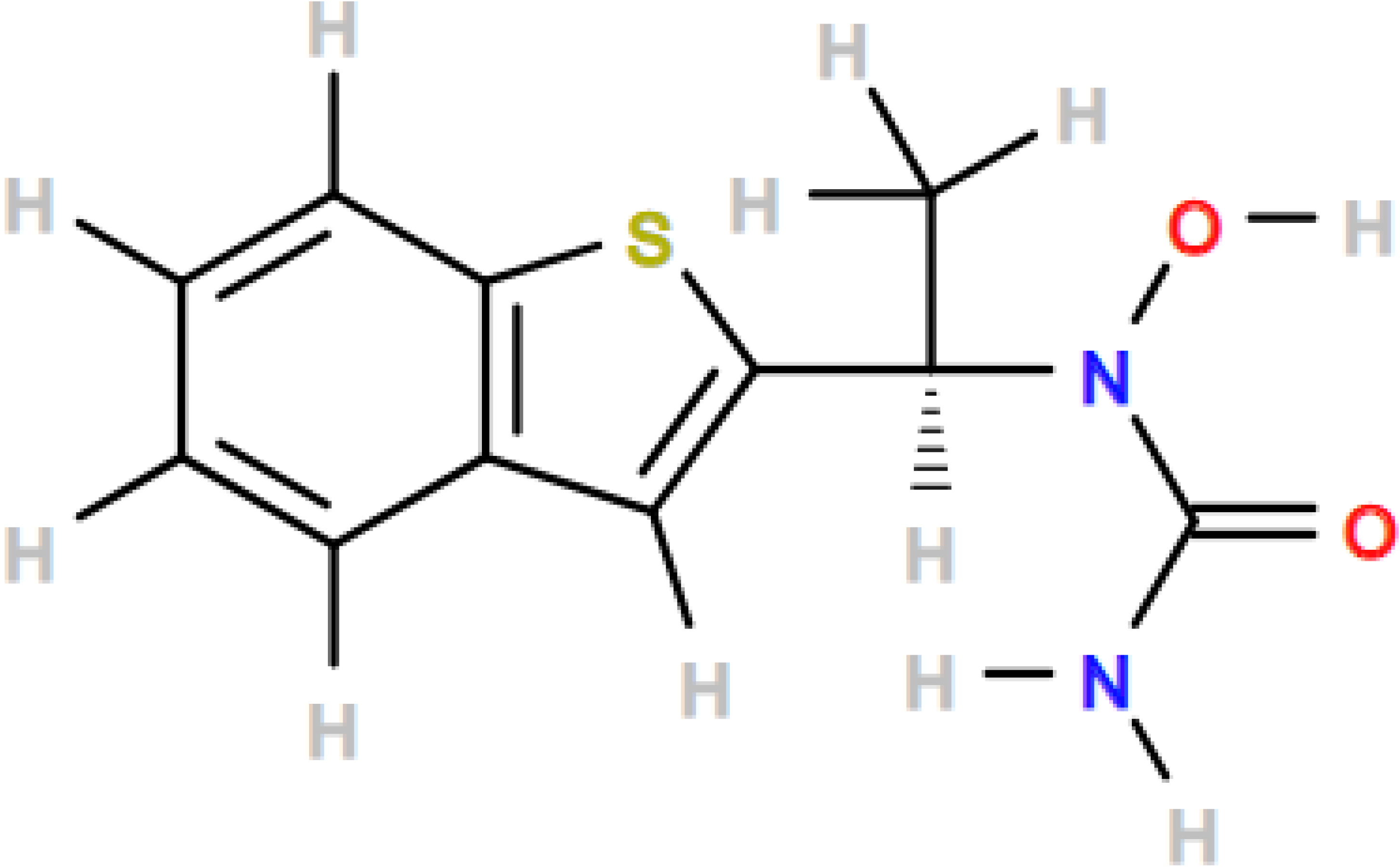

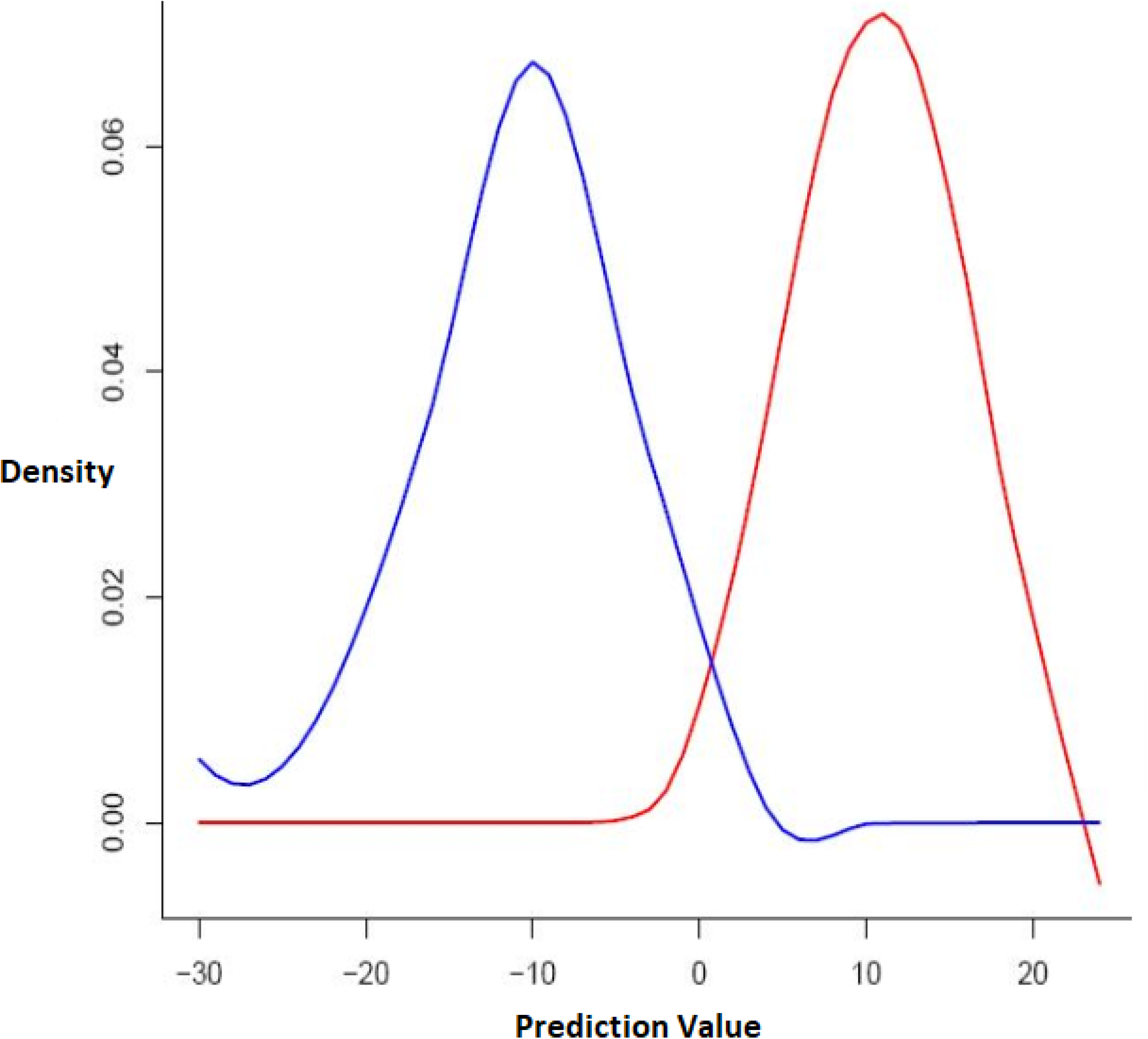

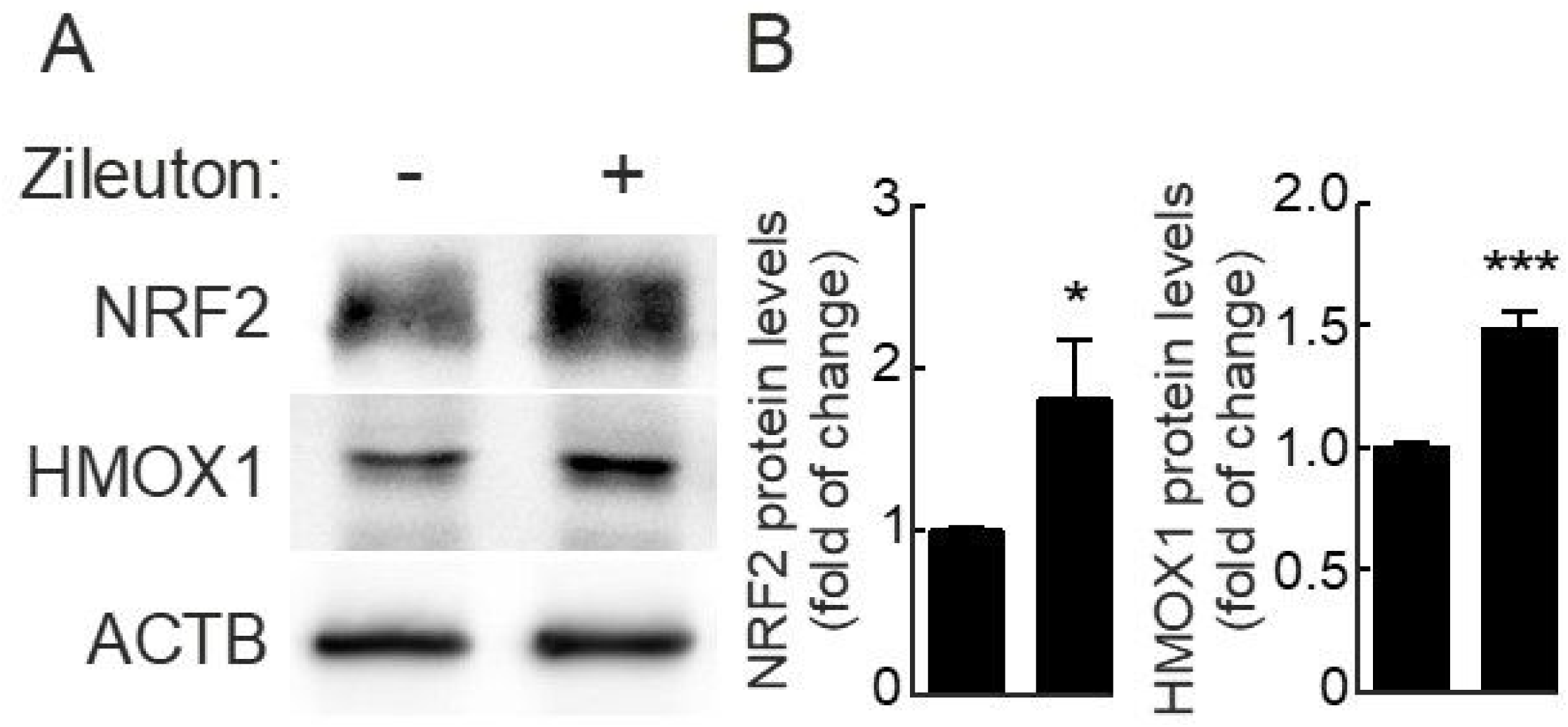

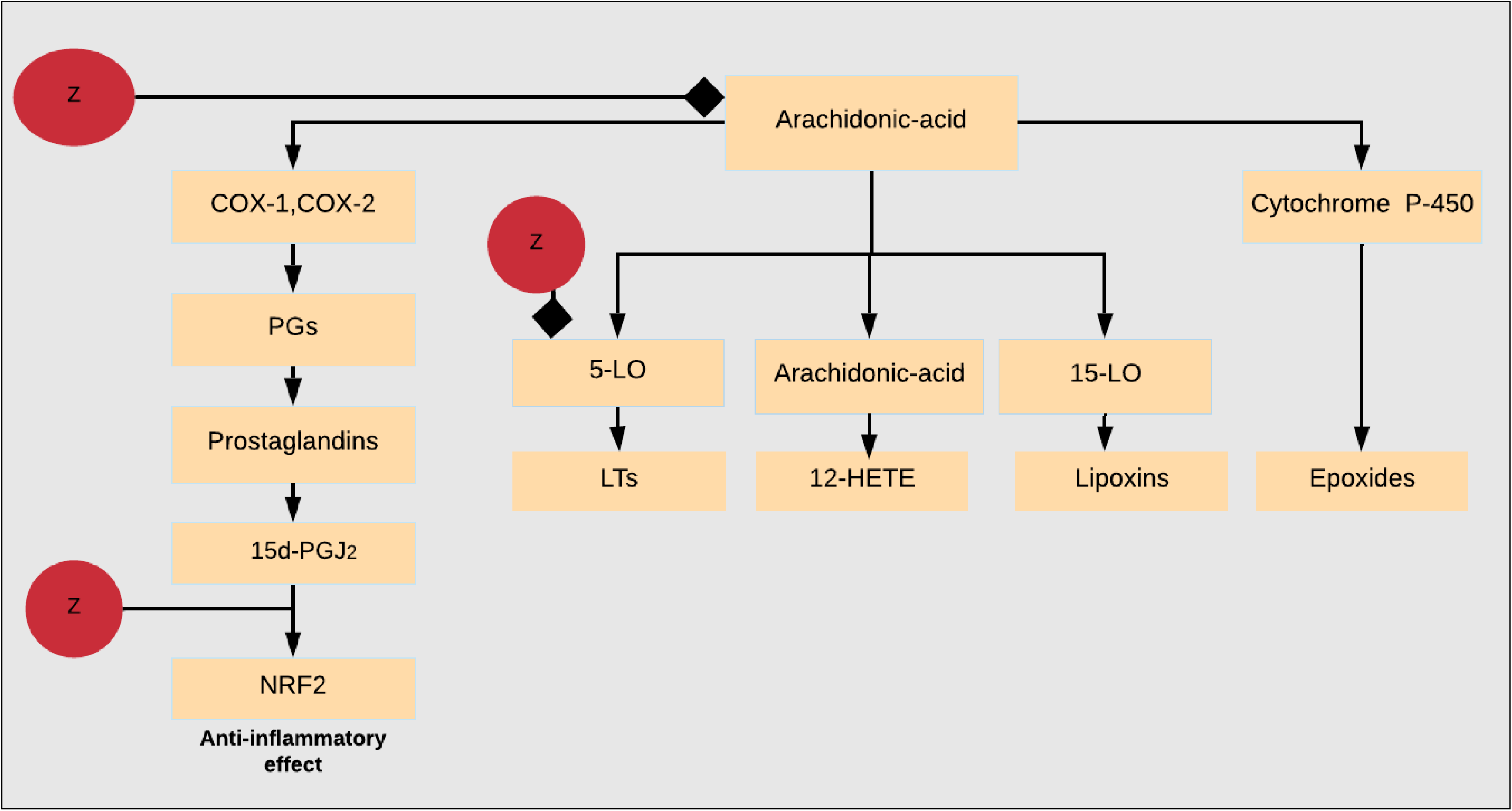

