## Supplementary material for "NRF2 drug repurposing using a question-answer artificial intelligence system": figures Legend

Figure 1. The workflow of the QAAI system consist of two pathways. PubMed articles are downloaded, parsed and tokenized, in the first pathway (ii) the second pathway covers the AI encoding of the query in relationship with the available dataset. Sentence embedding is followed by calculating a correlation matrix that identifies the right answer. The answers are filtered and then highly probable compounds ae tested for their ability to cross the blood brain barrier. Finally, experimental validation takes place.

Figure 2. Vector embedding. Each sentence in the document is converted to a unique vector of numbers using AI google semantic encoder.

Figure 3. Correlation matrix. A correlation matrix is produced between the question and all the parsed sentences available in the database. In this example, the questions posed is “what are the drugs that affect oxidative stress in chronic diseases?". The pipeline calculates sentence similarity for each sentence in the database.

Figure 4. F-score. Using a correlation matrix for estimating the right answer seems to be better than using a cosine matrix.

Figure 5. Zileuton chemical structure. Its molecular formula is C_11_H_12_N_2_O_2_S and it has molecular weight of 236.29.

Figure 6. Our workflow predict that Zileuton has the ability of crossing the BBB with score of 14.52 using AdaBoost algorithm.

Figure 7. Zileuton increases NRF2 levels. A, Representative immunoblots for the indicated proteins in cell lysates from RAW264.7 cells treated with zileuton (10 µM, 16 h) in the absence of serum. B, Densitometric quantification of representative immunoblots from A relative to ACTB protein levels. Data are mean ± SEM (n=4). Statistical analysis was performed using Student’s t test. *p<0.05; ****p<0.001 vs. vehicle-treated cells.

Figure 8. Zileuton model of action. In response to ROS stress, AA is released from membrane phopholipids by phospholipases. Free AA can be converted to bioactive eicosanoids through the cyclooxygenase (COX), lipoxygenase (LOX) or P-450 epoxygenase pathways. LOX enzymes (5-LO, 12-LO, 15-LO) catalyse the formation of LTs, 12(S)hydroperoxyeicosatetraenoic acids and lipoxins (LXs), respectively. COX isozymes (constitutive COX-1 and inducible COX-2) catalyse the formation prostaglandin. The P-450 epoxygenase pathway catalyses the formation of hydroxyeicosatetraenoic acids (HETEs) and epoxides. Zileuton was shown to inhibit 5-LO as well as prostaglandin production through suppress prostaglandin biosynthesis by inhibition of arachidonic acid release in macrophages. Zileuton can also activate NRF2.
