## Supplementary material for "NRF2 drug repurposing using a question-answer artificial intelligence system": Main suppl-table-mkr

Mkr-Table

| Method | TP | TN | FP | FN |
| --- | --- | --- | --- | --- |
| Inner product | 5 | 0 | 2 | 3 |
| Directed_hausdorff | 1 | 0 | 6 | 3 |

$$Accuracy=\frac{TP+TN}{TP+FP+FN+TN} (1)$$

$Precision=\frac{\mathrm{TP}}{TP+FP}$	(2)

$$Recall=\frac{\mathrm{TP}}{TP+FN} (3)$$

$F_{1}^{\mathrm{Score}}=2*\frac{Precision * Recall}{Precision + Recall}$	(4)
